# Type-2 diabetes with low LDL-C: genetic insights into a unique phenotype

**DOI:** 10.1101/837013

**Authors:** Yann C. Klimentidis, Amit Arora, Michelle Newell, Jin Zhou, Jose M. Ordovas, Benjamin J. Renquist, Alexis C. Wood

## Abstract

Although hyperlipidemia is traditionally considered a risk factor for type-2 diabetes (T2D), evidence has emerged from statin trials and candidate gene investigations suggesting that lower LDL-C increases T2D risk. We thus sought to comprehensively examine the phenotypic and genotypic relationships of LDL-C with T2D. Using data from the UK Biobank, we found that LDL-C was negatively associated with T2D (OR=0.43[0.41, 0.45] per mmol/L unit of LDL-C), despite positive associations of LDL-C with HbA1c and BMI. We then performed the first genome-wide exploration of variants simultaneously associated with lower LDL-C and increased T2D risk, using data on LDL-C from the UK Biobank (n=431,167) and the GLGC consortium (n=188,577), and T2D from the DIAGRAM consortium (n=898,130). We identified 31 loci associated with lower LDL-C and increased T2D, capturing several potential mechanisms. Seven of these loci have previously been identified for this phenotype, and 9 have previously been implicated in non-alcoholic fatty liver disease. Finally, two-sample Mendelian randomization analyses suggest that low LDL-C causes T2D, although causal interpretations are challenging due to pleiotropy. Our findings extend our current understanding of the higher T2D risk among individuals with low LDL-C, and of the underlying mechanisms, including those underlying the diabetogenic effect of LDL-C-lowering medications.

## Introduction

Rates of cardiovascular disease (CVD) and type-2 diabetes (T2D) are among the most pressing health concerns worldwide. These two diseases share many risk factors, and tend to co-occur; T2D carries a 2-4 fold increase in risk for CVD, and more than 70% of patients with T2D will die from cardiovascular complications (1). Yet, there remains controversy over whether all risk factors exert similar effects on the risk of these two conditions. Low-density lipoprotein cholesterol (LDL-C) is a class of highly atherogenic particles, and circulating levels of LDL-C are a causal risk factor for CVD across the lifespan (2). Lipid lowering medications, in particular from the statin drug class, are effective at lowering levels of LDL-C which has a dose-response relationship with a reduction in adverse cardiovascular events (3). The perception of a strong inter-relationship of T2D with hyperlipidemia, in particular higher LDL-C, has led to a standard of care in which lipid-lowering medications are a common therapeutic option for T2D. For example, the American Diabetes Association (ADA) recommends high intensity statin use for individuals with diabetes with prevalent atherosclerotic cardiovascular disease (ASCVD) or a greater than 20% 10-year risk of ASCVD until they reach recommended stringent targets of LDL-C < 70mg/dL. The ADA further recommends moderate-intensity statin use for T2D patients 40 years and older without ASCVD (4).

A recent analysis of national registry data from Scotland showed that 83.6% and 71.1% of T2D patients with and without CVD were on a statin, respectively (5). Similar figures have emerged from analyses of Norwegian (6) and US populations (7). This almost ubiquitous use of statins in T2D is highly concerning given that several lines of evidence suggest that lowered LDL-C is a causal risk factor for T2D. A small number of observational studies have reported that individuals with low levels of LDL-C (e.g. <60 mg/dl), exhibit a higher risk of T2D (8,9), and that among individuals with coronary disease, LDL-C and T2D are inversely related (10). Further, individuals with familial hypercholesterolemia exhibit a decreased risk of T2D (11). Clinical trials have found that statin use conveys an increased T2D risk (odds ratio=1.09) (12,13) in a dose-dependent manner (14). Although statins do effectively lower LDL-C, LDL is key to delivering hepatic lipids from the liver to peripheral tissues. Accordingly, by lowering LDL-C production, statins mute the ability of the liver to export lipids. Given that the incidence and severity of hyperinsulinemia and insulin resistance is directly related to hepatic lipid concentrations, it is easy to understand how limiting hepatic lipid export may indirectly increase the incidence of T2D (15,16).

Genetic studies have lent further support to inverse phenotypic associations between LDL-C and T2D, with recent studies pointing to genetic loci that harbor variants exerting opposing effects on LDL-C and T2D. These include loci containing the *HMGCR* (17), *APOE* (18,19), *PCSK9* (17,19,20), *NPC1L1* (17,19), *PNPLA3* (19), *TM6SF2* (19), *GCKR* (19), and *HNF4A* (19) genes. Furthermore, both Fall et al. (21) and White et al. (22) have found that genetically-predicted higher LDL-C was associated with a lower risk of T2D. Yet, genetic findings show that not all variants have opposing effects on LDL-C levels and T2D risk. LDL-C lowering variants in *ABCG5/G8* and *LDLR* genes were not shown to alter T2D risk (17) and subsets of LDL-C lowering alleles provide a stronger prediction against T2D than the full gamut (21). LDL-C levels, like T2D, are reflective of a number of physiological processes. The findings outlined above suggest that there is heterogeneity in T2D outcomes, depending on which pathways are the primary LDL-C lowering mechanisms, and that genetic studies may be able to give us insights into these pathways. A better understanding of which genetic loci lower LDL-C but increase T2D risk may thus yield mechanistic insights that may help develop therapeutic options that lower lipid levels without raising the risk of T2D, and help identify individuals at greater risk for T2D with statin use.

The conclusion of the ADA’s Professional Practice committee is that “the cardiovascular event rate reduction with statins far outweighed the risk of incident diabetes” due to their belief that the increase in risk “may be limited to those with diabetes risk factors”. By studying genetic loci that are inversely associated (or not) with LDL-C and T2D risk, we aimed to identify pathways and therapeutic targets that may be harnessed to develop optimal therapeutic options.

Here, we first examined the relationship of directly-measured LDL-C with prevalent T2D, HbA1c, and BMI. We then sought to identify, for the first time on a genome-wide scale, loci simultaneously associated with lower LDL-C and increased T2D (and vice versa). Upon identifying variants, we sought to generate additional mechanistic insights by testing of associations with seven other traits in the UK Biobank related to T2D, LDL-C, and non-alcoholic fatty liver disease (NAFLD). Finally, we performed two-sample Mendelian randomization (MR) analyses to examine a potential causal effect of LDL-C on risk of T2D.

## Methods

### UK Biobank

Data from the UK Biobank was used for (1) phenotypic data analysis, which examined the associations of LDL-C and TG with T2D, HbA1c, and BMI, (2) discovery GWAS for variants which are associated with lower LDL-C and higher T2D, and (3) creation of a LDL-C genetic instrument to test in a 2-sample MR framework for a potential causal relationship between LDL-C and T2D. The UK Biobank is a prospective cohort study of approximately 500,000 individuals between the ages of 39 and 72, living throughout the United Kingdom (UK). Participants attended one of 21 assessment centers in the UK and had their blood drawn for biomarker and genetic analysis, and weight and height measured to derive BMI (kg/m^2^). Directly-measured LDL-C, HbA1c, HDL, TG, alanine aminotransferase (ALT) and aspartate aminotransferase (AST) were obtained from all UK Biobank participants at the baseline visit between 2006 and 2010 in a non-fasting state. LDL-C was assessed by enzymatic protective selection analysis on a Beckman Coulter AU5800. Prevalent T2D was defined using the following criteria: 1) self-reported T2D or generic diabetes in verbal interviews, 2) over 35 years at age of diagnosis, and 3) not using insulin within one year of diagnosis, to exclude possible type-1 diabetes cases (23).

### UK Biobank genotypes

Genotypes in the UK Biobank were obtained with the Affymetrix UK Biobank Axiom Array (Santa Clara, CA, USA), while 10% of participants were genotyped with the Affymetrix UK BiLEVE Axiom Array. Details regarding imputation, principal components analysis, and QC procedures are described elsewhere (24). Individuals with unusually high heterozygosity, with a high (>5%) missing rate, or with a mismatch between self-reported and genetically-inferred sex were excluded from analyses. SNPs out of Hardy-Weinberg equilibrium (p<1 x10^−6^), with a high missing rate (>1.5%), with a low minor allele frequency (<0.1%), or with a low imputation accuracy (info<0.4) were excluded from analyses. This resulted in the availability of approximately 15 million SNPs for analysis.

### DIAGRAM and GLGC GWAS meta-analysis summary statistics

The latest GWAS meta-analysis summary statistics for T2D (unadjusted for BMI) were obtained from the DIAGRAM (Diabetes Genetics Replication and Meta-Analysis) consortium that includes data on up to 898,130 individuals (74,124 and 824,006 controls), including UK Biobank individuals (25). We used the results of our GWAS of LDL-C in UKB along with the aforementioned DIAGRAM-T2D results for the discovery of inverse association signals. We then replicated LDL-C associations of our top hits with an independent GWAS meta-analysis of LDL-C from the GLGC (26) (Global Lipids Genetics Consortium; n=188,577; this meta-analysis does not include the UK Biobank study). Across the UK Biobank, DIAGRAM, and GLGC summary statistics, we aligned all SNP alleles and their corresponding effects by using the harmonize function in the TwoSampleMR package in R (27).

### Mendelian randomization

In a two-sample Mendelian randomization (MR) framework, we used 185 independent variants associated with LDL-C in the UK Biobank (as detailed below) as the genetic exposure, and the results of a previous DIAGRAM T2D GWAS meta-analysis (does not include the UK Biobank) as the outcome (28). The TwoSampleMR package in R was used to perform analyses. We used the *clump_data* function to identify independent exposure SNPs (r^2^ < 0.001 in 1,000 Genomes Project European samples) from all SNPs with p<5 × 10^−8^, and excluded unresolvable strand-ambiguous SNPs and any SNPs for which no proxy could be identified (r^2^<0.8). We examined results from the inverse-variance weighted (IVW), MR-Egger, and weighted median analyses.

### Statistical analyses

To evaluate and plot the prevalence of T2D, and of BMI and HbA1c, by decile of LDL-C and TG in the UK Biobank, we excluded all participants who self-reported (at baseline) use of cholesterol-lowering medications during the touchscreen survey, or statin medication during the verbal interview. To examine levels of HbA1c by decile of LDL-C, we excluded participants who self-reported T2D at the baseline visit. We further excluded individuals with outlier values of HbA1c greater than four standard deviations from the mean. Deciles were calculated using the ‘quantcut’ function in the ‘gtools v3.5.0’ library in R. Once deciles were established, T2D prevalence by LDL-C/TG deciles was calculated and plotted with confidence intervals determined by the Clopper-Pearson interval (29). Mean HbA1c and BMI and their distributions are shown in boxplots for each decile of LDL-C. We further examined T2D prevalence by LDL-C decile separately in males and females, and in different age groups (40-49 years, 50-59 years, and 60-69 years).

To statistically evaluate these phenotypic associations, we performed logistic regression with T2D as the outcome, and linear regression with HbA1c and BMI as outcomes. For each analysis, we used the same exclusion criteria as those mentioned above, and adjusted for ‘last eating’ time (excluding individuals reporting extreme values, >16 hours), age, sex, and center. To normalize residuals, we transformed LDL-C, TG cholesterol, HbA1c, and BMI by inverse normalization for all linear regression analyses. In analyzing the association of LDL-C with T2D, we also tested for interactions with sex and age and provided stratified analyses accordingly.

For the genome-wide association study of LDL-C in the UK Biobank, the LDL-C level of individuals on cholesterol-lowering medication was corrected by dividing it by a correction factor of 0.63 (30). We transformed LDL-C by inverse normalization. We used BOLT-LMM software (31) to perform GWAS on individuals of European descent (n=431,167) and included ‘last eating’ time (see above), sex, age, age^2^, center, genotyping chip and the first 10 PCs as covariates. BOLT-LMM performs a linear mixed model regression that includes a random effect of all SNP genotypes other than the one being tested. We aligned effect sizes across the GWAS summary statistics of each trait to the same effect allele using the *harmonize* function, as mentioned above. We then selected only those SNPs that exhibited opposite directions of effects for LDL-C and T2D. Among these, we selected only those with an association p-value < 5 × 10^−5^ for LDL-C and for T2D in a region defined as a maximum size of 500 kb. At this p-value threshold, the prior probability of a given SNP associated with two traits and with discordant direction of effect under the null hypothesis, corresponds to 0.00005 × 0.000025 = 1.25 × 10^−9^ (32). We did not employ a multivariate approach because current methods do not typically report trait-specific effect size estimates (or directions). Although MTAG (33) does provide effect size estimates for each trait, this method is not suitable for traits with low genetic correlation, which is the case for LDL-C and T2D (25). For the replication of the discovered loci, we used a threshold of p<5 × 10^−3^ in GLGC (does not include UK Biobank individuals), and exhibiting the opposite direction of association with T2D in DIAGRAM. To test the association of the 31 SNPs (T2D increasing allele) that we identified with a range of other cardiometabolic traits, we used similar methods described above for the LDL-C GWAS. For TG and HDL-C, we excluded individuals reporting cholesterol-lowering medication. For ALT and AST, we excluded 15,138 individuals with medical conditions, other than NAFLD, that could affect liver enzyme levels (34). For HbA1c, we excluded individuals with self-reported T2D (see above). For WHR, we additionally adjusted for BMI prior to inverse normalization and subsequent GWAS. We inverse normalized all traits before GWAS. After association of 31 SNPs with each of these seven additional phenotypes, we normalized the effect sizes by dividing the beta coefficients by the corresponding standard errors, and dividing by the square root of the respective sample size. We then used hierarchical clustering to group the identified variants according to their pattern of association with all nine traits, including T2D and LDL-C. We used the *hclust* function in R, with the Euclidian metric to calculate distances, and the Ward clustering method (35).

## Results

### Participant characteristics

In a sample size of 379,617 individuals after exclusions of individuals on lipid-lowering medication, T2D prevalence was 0.9%, and was higher in males (1.53 %) than females (0.63 %). Individuals with T2D had lower LDL-C, higher TG, higher HbA1c, and higher BMI (see Supplementary Table 1).

### Association of LDL-C with T2D

We observed an inverse relationship between LDL-C and T2D prevalence (T2D prevalence = 0.90%; OR=0.43 [0.41, 0.45], p=2.6 × 10^−269^). Individuals in the lowest decile of LDL-C exhibited the highest prevalence of T2D and a consistent decrease in T2D prevalence was observed with increasing LDL-C (see Figure 1). We found a very similar negative association of LDL-C with T2D among only the individuals reporting the use of cholesterol-lowering medication. We found a significant interaction of LDL-C with sex (p=1.2 × 10^−18^), whereby the association of LDL-C with T2D was stronger among men (OR=0.36 [0.34, 0.38], p=3.6 × 10^−227^) than among women (OR=0.54 [0.50, 0.58], p=3.63 × 10^−60^) (see Supplementary Table 2 and Supplementary Figure 1). We also observed a stronger inverse association between LDL-C and T2D prevalence among older individuals (Supplementary Table 2 and Supplementary Figure 2). Positive associations were found between LDL-C and both HbA1c (after exclusion of individuals with T2D; β = 0.14, se = 0.0017, p < 2.2 × 10^−308^) and BMI (β = 0.16, se = 0.0016, p< 2.2 × 10^−308^) (Figure 1 and Supplementary Table 2). We also observed a positive association between TG and T2D prevalence (OR=1.35 [1.32, 1.39], p=2.4 × 10^−134^; Supplementary Table 2 and Supplementary Figure 3).

**Figure 1:**
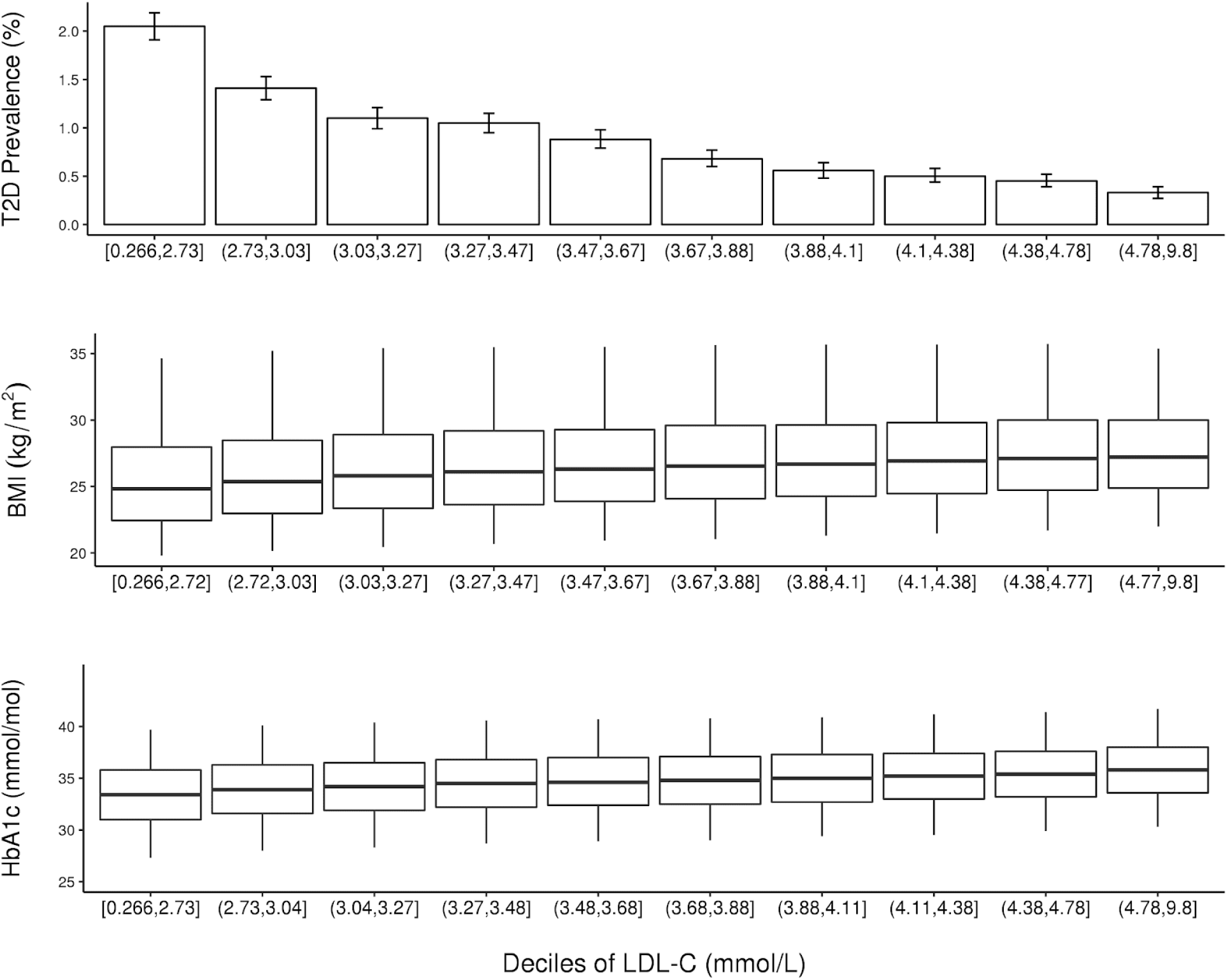
T2D prevalence, HbA1c, and BMI by LDL-C decile in the UK Biobank. T2D prevalence is shown as a percentage with error bars corresponding to the Clopper-Pearson confidence interval. Whisker plots show the median value (horizontal line in box), the 25^th^ and 75^th^ percentile delimited by the box, and the vertical lines extending to the 5^th^ and 95^th^ percentile.

### Loci associated inversely with LDL-C and T2D

We identified 48 loci associated in opposite directions with LDL-C and T2D using the UK Biobank LDL-C and the DIAGRAM T2D results. Among these, 31 replicated with respect to LDL-C association when using the GLGC LDL-C GWAS results instead of UK Biobank (see Table 1). Several loci are previously known or suspected to be inversely associated with LDL-C and T2D (*HMGCR, APOE, NPC1L1, PNPLA3, TM6SF2, GCKR*, and *HNF4A*). However, most of the loci are novel for this LDL-T2D trait. Of these novel loci, 12 have previously been identified for LDL-C in the GLGC GWAS, and 14 were previously identified in T2D GWAS, and 14 have not been identified previously with either trait. The loci with the strongest degree of opposing effects include *FNDC7-STXBP3, SORT1-PSMA5, HMGCR-POC5, PPP1R3B*, and *GCKR* (see Figure 2).

**Table 1:**
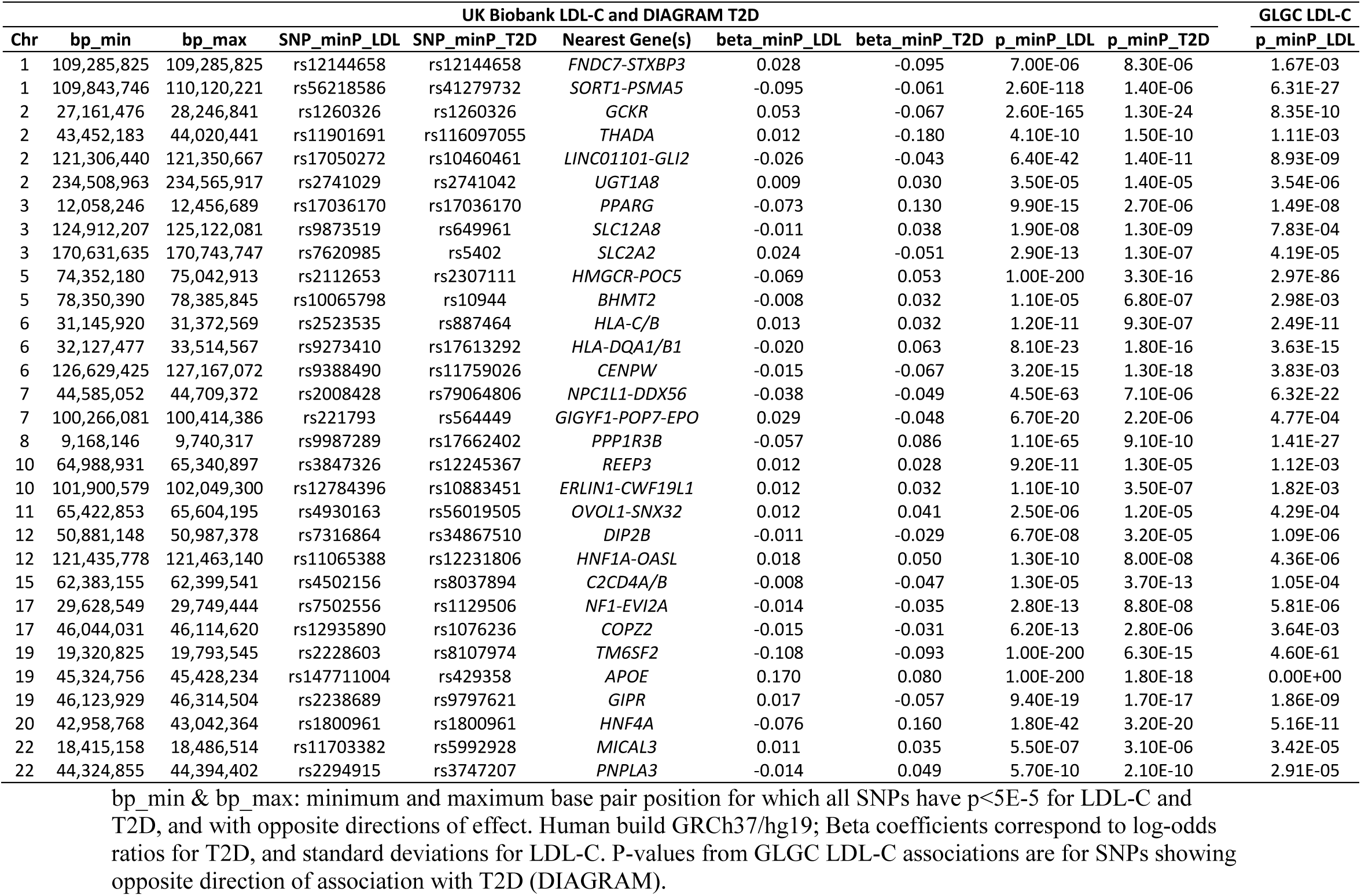
List of genetic loci in which all SNPs have p<5E-5 for both LDL-C and T2D and with opposite directions of effect, for UK Biobank LDL-C and DIAGRAM T2D. Beta coefficients refer to the SNP with the lowest p-value for each trait in the respective region. The minimum p-value for LDL-C in GLGC summary statistics for the region discovered in UK Biobank is also shown.

**Figure 2:**
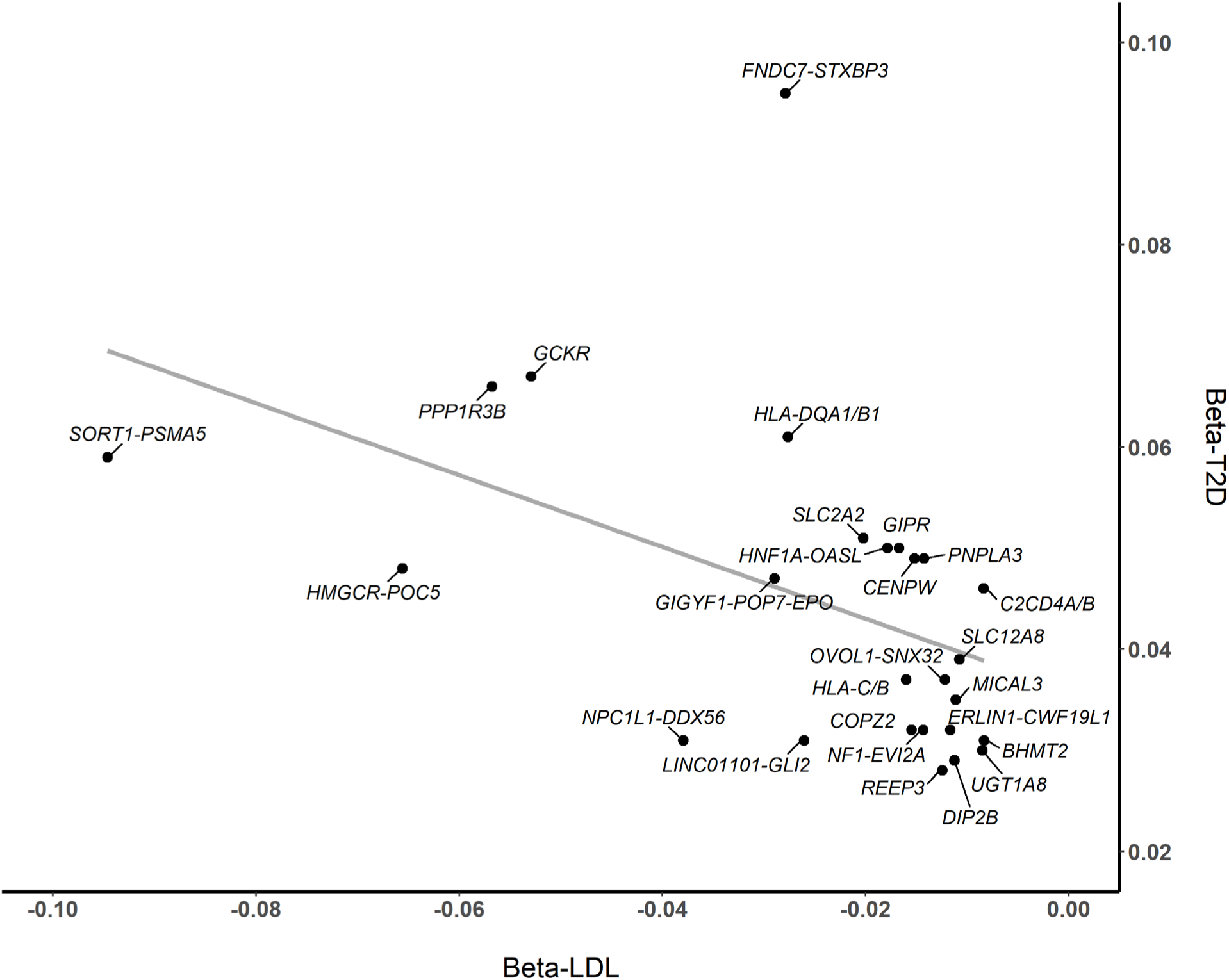
Plot of beta coefficients for LDL-C vs. T2D for SNPs with opposite directions of effect on these two traits. Beta coefficients correspond to log-odds ratios for T2D, and standard deviations for LDL-C.

The variants that we have identified can be linked with genes that affect *de novo* fatty acid synthesis, hepatic lipid uptake, hepatic lipid export, peripheral tissue lipid balance, fatty liver of unknown origin, insulin secretion, and insulin action (see Supplementary Table 4). They are associated in distinct patterns across a range of cardiometabolic traits (see Figure 3). At these loci, the T2D-increasing alleles at the SNPs with lowest T2D p-value are generally associated with higher HbA1c levels, lower HDL levels, and a higher WHR, although this pattern is not entirely consistent across all 31 SNPs. A cluster of *APOE, GCKR*, and *TM6SF2* emerged showing a pattern of lower TG, higher HbA1c, and generally higher BMI and WHR. Another cluster includes *ERLIN1-CWF19L1* and *PNPLA3* that exhibits lower HDL, and higher liver enzymes. Finally, another cluster emerged including *SORT1-PSMA5, HMGCR-POC5*, and *HNF4A*, which generally exhibit higher BMI and lower HDL (see Figure 3).

**Figure 3:**
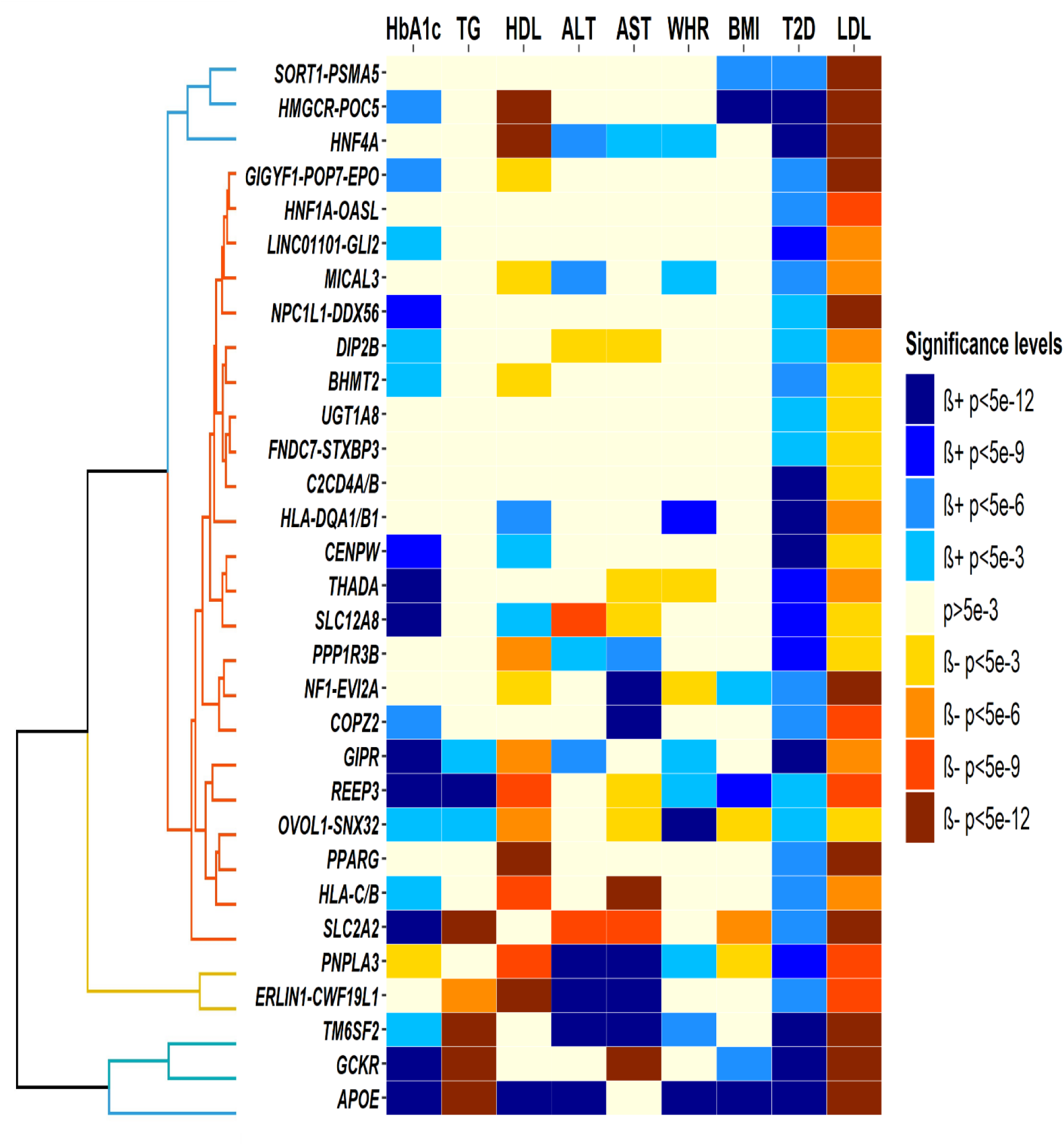
Association of T2D increasing allele at 31 identified SNPs with nine cardiometabolic traits, and clustering dendogram based on corresponding standardized effect sizes.

### Mendelian randomization

We used 140 independent SNPs significantly associated with LDL-C in the UK Biobank as the genetic instrument. We found a consistent negative association with T2D across all MR methods implemented (IVW OR=0.82 [0.72, 0.93]) except for simple mode, although we noted a high degree of heterogeneity (see Table 2 and Supplementary Figures 4-6).

**Table 2:** Results of Mendelian randomization of LDL-C on type-2 diabetes.

| Method | n SNP | OR [95% CI] | p-value | Modified<br>Cochran's Q,<br>p-value | MR-Egger<br>Y-intercept,<br>p-value |
| --- | --- | --- | --- | --- | --- |
| MR Egger | 140 | 0.68 [0.54, 0.85] | 8.58E-04 | 1.01E-04 | 0.05 |
| Weighted median | 140 | 0.79 [0.66, 0.96] | 1.98E-02 |  |  |
| Inverse variance weighted | 140 | 0.82 [0.72, 0.93] | 2.64E-03 | 4.02E-05 |  |
| Simple mode | 140 | 0.79 [0.52, 1.21] | 2.80E-01 |  |  |
| Weighted mode | 140 | 0.81 [0.68, 0.96] | 1.82E-02 |  |  |

## Discussion

We used the largest sample to date to demonstrate that LDL-C is inversely correlated with T2D risk, such that individuals with low LDL-C exhibit a higher prevalence of T2D. Then, in the first genome-wide analysis aimed at identifying variants associated with both lower LDL-C and higher T2D risk, we identified 24 novel loci exerting opposite-direction effects on these traits. Our analyses lend weight to the notion that the association between lower LDL-C and increased T2D risk is driven, at least in part, by a specific group of genetic variants that may be implicated via diverse mechanisms including hepatic lipid synthesis, export, and uptake, as well as insulin secretion and action. These variants provide insight into the heterogeneous outcomes for different lipid and glucose metabolism pathways and may also point to novel susceptibility loci for hyperlipidemia and T2D.

We find that low LDL-C is associated with greater T2D risk, which is consistent with two previous studies examining T2D prevalence (8) and incidence (9). In addition, we find that lower LDL-C is associated with lower HbA1c (among individuals without T2D) and lower BMI. On the other hand, we do observe a similar inverse relationship of LDL-C with T2D in the set of people on cholesterol-lowering medication. We also find that unlike LDL-C, TG levels are positively associated with T2D prevalence. This opposing relationship of LDL-C and TG with T2D suggests that LDL are being overfilled in individuals with T2D.

Previous research into loci that jointly alter the risk for LDL-C and T2D has focused on the genomic targets of lipid-lowering medications, in the hope that these analyses will give specific insights into associated T2D risk. Our analyses confirmed that variants in *HMGCR* (36) and *NPC1L1* (19) are associated with lower LDL-C and increased T2D risk. On the other hand, our analyses did not identify variants at *PCSK9*. The lowest T2D p-value was 0.003 in this region for a SNP with opposite direction coefficient. However, our analyses identified a fourth target of lipid-lowering medications: variants in the Peroxisome Proliferator-Activated Receptor (*PPARG*) gene, the target of fibrates and thiazolidinediones.

We observed nine variants previously identified as being associated with NAFLD: *PNPLA3, GCKR, TM6SF2, PPP1R3B, ERLIN1*-*CWF19L1, REEP3, HNF1A, SLC2A2*, and *MICAL3* (37–39). This enrichment for NAFLD-related genes may reflect increased synthesis and storage of TG and reduced export/secretion of VLDL, leading to reduced circulating LDL-C. Indeed, the LDL-C-decreasing alleles at most of these loci are associated with increased liver enzymes, indicative of hepatic steatosis (with the exception of *GCKR* and *SLC2A2*, consistent with a previous finding (40)). The degree of NAFLD is directly linked to the incidence and severity of hyperinsulinemia and insulin resistance in obesity (15,16). In turn, lower LDL-C with increased liver enzymes would be expected to indicate increased NAFLD and T2D. A recent bi-directional Mendelian randomization study provides support for this hypothesized causal effect of NAFLD on T2D (41). Our findings that liver fat may be an important mediator of the effect of cholesterol-lowering on T2D is consistent with a report showing that liver fat may help identify statin-taking individuals at risk of T2D (42). Finally, it is noteworthy that the *HMGCR* variant that lowers LDL-C is not associated with any significant change in liver enzymes, potentially reflecting the lack of an increase in NAFLD incidence seen with statin medications (43).

Our analyses identified a number of variants previously implicated in lipid and glucose metabolism. Sortilin 1 (*SORT1*) is highly expressed in adipocytes, and the sortilin gene product facilitates the formation and export of VLDL from the liver (44,45). The role of *SORT1* in T2D risk is not well understood. Sortilin 1 is required for insulin-dependent glucose uptake (46–48), yet *Sort1* knockout mice may show reduced glucose and glycolic intermediates in the fasted state (49). This highlights again the potential for heterogeneous paths in T2D risk and the dependence on multiple pathways of lipid and glucose metabolism to explain our findings.

Several loci were also identified which are known to be related to T2D, without known associations with LDL-C. These include *THADA, C2CD4A, CENPW*, and *SLC12A8* (25). In addition, we identified several variants associated with lower LDL-C but increased T2D risk with no known biological pathways linking these loci to either trait. *SLC2A2*, which encodes the glucose transporter 2, has not previously been associated with either LDL-C or T2D in the large respective GWAS consortia. However, GLUT2 is key to hepatic glucose uptake following a meal and the associated hepatic *de novo* lipogenesis (50). In fact, liver-specific GLUT2 knockout decreases liver triglyceride concentrations. Importantly, GLUT2 expression in the β-cell is required for the glucose stimulated insulin response (51). In turn, a locus that decreases GLUT2 expression would be expected to limit serum insulin, increase HbA1c, and decrease LDL-C.

Our approach is subject to several limitations. We used prevalent T2D in UK Biobank, which limits inferences related to the direction of causality. As incident T2D cases develop in the UK Biobank, it will be important to examine the association of LDL-C at baseline with incident T2D. It is also difficult to identify the causal gene at identified loci. Although we annotated these loci according to nearby genes and/or previous annotation, the listed genes may not necessarily be directly implicated, if at all. Finally, it is difficult to draw any firm conclusions from MR as these traits are highly intertwined and the assumption of no pleiotropy is unrealistic.

In conclusion, our results suggest that low-LDL-C may be a risk factor for T2D, although further study is warranted. We have identified a collection of genetic variants that may provide insight into the mechanisms underlying the diabetogenic risk of low LDL-C, and of lipid-lowering medications, and the decreased T2D risk among individuals with familial hypercholesterolemia.

## Acknowledgments

The authors would like to acknowledge the vital contributions of the GLGC and DIAGRAM consortia, as well as all organizers and participants of individual participating studies. This research was conducted using the UK Biobank Resource under Application Number 15678. We thank the participants and organizers of the UK Biobank. The authors would like to acknowledge support from the National Heart, Lung, and Blood Institutes (R01-HL136528). Dr. Wood’s role on this project was funded, in part, by USDA/ARS cooperative agreement # 58-3092-5-001. The contents of this publication do not necessarily reflect the views or policies of the U.S. Department of Agriculture, nor does mention of trade names, commercial products, or organizations imply endorsement by the U.S. Government. The funders had no role in study design, data collection and analysis, decision to publish, or preparation of the manuscript.

The authors have no relevant conflicts of interest to disclose.

## Author Contributions

YK conceived and designed the study. AA, MN, and YK performed data analyses. YK, AW, MN, and BR wrote the manuscript. AW, BR, and JO contributed to writing of the introduction and discussion; all authors read and edited the manuscript.

