## Supplemental Material for "Type-2 diabetes with low LDL-C: genetic insights into a unique phenotype"

**Supplementary Table 1:** Descriptive statistics for UK Biobank participants (N = 379,617) included in LDL-C vs. T2D analysis, presented by T2D prevalence.

|  | **Prevalent T2D** | |
| --- | --- | --- |
|  | **Yes** | **No** |
| **N** | 3,416 | 376,201 |
| **Age ± sd (years)** | 58.95 ± 7.26 | 55.47 ± 8.08 |
| **Sex (% Female)** | 40.19% | 57.88% |
| **LDL ± sd (mmol/L)** | 3.29 ± 0.77 | 3.72 ± 0.81 |
| **TG ± sd (mmol/L)** | 2.20 ± 1.25 | 1.69 ± 1.00 |
| **HbA1c ± sd (mmol/mol)** | 49.98 ± 13.28 | 34.76 ± 3.76 |
| **BMI ± sd (kg/m^2^)** | 31.39 ± 6.11 | 26.94 ± 4.58 |

**Supplementary Table 2:** Regression analysis for T2D, HbA1c, and BMI with LDL-C and TG.

| **LDL-C** |  |  | |  | |  | |
| --- | --- | --- | --- | --- | --- | --- | --- |
|  | **Dependent variable (binary)** | **OR** | **95% CI** | | **p-value** | | **N** |
|  | T2D | 0.43 | [0.41, 0.45] | | 2.63E-269 | | 377,418 |
|  | T2D, LDL-C*sex | 0.65 | [0.59, 0.72] | | 1.22E-18 | | 377,418 |
|  | T2D, LDL-C*age | 0.97 | [0.97, 0.98] | | 3.03E-18 | | 377,418 |
|  | T2D, sex-female | 0.54 | [0.50, 0.58] | | 3.63E-60 | | 217,994 |
|  | T2D, sex-male | 0.36 | [0.34, 0.38] | | 3.55E-227 | | 159,424 |
|  | T2D, ages 40-49 years | 0.71 | [0.63, 0.81] | | 1.26E-07 | | 102,347 |
|  | T2D, ages 50-59 years | 0.43 | [0.40, 0.47] | | 2.74E-87 | | 132,657 |
|  | T2D, ages 60-69 years | 0.37 | [0.35, 0.40] | | 1.11E-188 | | 141,029 |
| **LDL-C** |  |  |  | |  | |  |
|  | **Dependent variable (continuous)** | **Beta** | **SE** | | **p-value** | | **N** |
|  | HbA1c | 0.14 | 1.65E-03 | | < 2.23E-308 | | 353,597 |
|  | BMI | 0.16 | 1.63E-03 | | < 2.23E-308 | | 377,214 |
| **TG** |  |  |  | |  | |  |
|  | **Dependent variable (binary)** | **OR** | **95% CI** | | **p-value** | | **N** |
|  | T2D | 1.35 | [1.32, 1.39] | | 2.40E-134 | | 377831 |

**Supplementary Table 3:** List of loci identified in discovery analysis with UKB LDL-C and DIAGRAM T2D.

| **Chr** | **bp_min** | **bp_max** | **SNP_minP_LDL** | **SNP_minP_T2D** | **Nearest Gene** | **beta_minP_LDL** | **beta_minP_T2D** | **p_minP_LDL** | **p_minP_T2D** |
| --- | --- | --- | --- | --- | --- | --- | --- | --- | --- |
| 1 | 109285825 | 109285825 | rs12144658 | rs12144658 | *FNDC7-STXBP3* | 0.0279078 | -0.095 | 7.00E-06 | 8.30E-06 |
| 1 | 109843746 | 110120221 | rs56218586 | rs41279732 | *SORT1-PSMA5* | -0.0946204 | -0.061 | 2.60E-118 | 1.40E-06 |
| 2 | 27161476 | 28246841 | rs1260326 | rs1260326 | *GCKR* | 0.0529385 | -0.067 | 2.60E-165 | 1.30E-24 |
| 2 | 43452183 | 44020441 | rs11901691 | rs116097055 | *THADA* | 0.0120544 | -0.18 | 4.10E-10 | 1.50E-10 |
| 2 | 121306440 | 121350667 | rs17050272 | rs10460461 | *LINC01101-GLI2* | -0.0260593 | -0.043 | 6.40E-42 | 1.40E-11 |
| 2 | 234508963 | 234565917 | rs2741029 | rs2741042 | *UGT1A8* | 0.00854497 | 0.03 | 3.50E-05 | 1.40E-05 |
| 3 | 12058246 | 12456689 | rs17036170 | rs17036170 | *PPARG* | -0.0727846 | 0.13 | 9.90E-15 | 2.70E-06 |
| 3* | 88263487 | 88264569 | rs7624215 | rs6419869 | *C3orf38-EPHA3* | 0.0164543 | -0.049 | 3.90E-08 | 1.10E-05 |
| 3 | 124912207 | 125122081 | rs9873519 | rs649961 | *SLC12A8* | -0.0107899 | 0.038 | 1.90E-08 | 1.30E-09 |
| 3* | 129300159 | 129301874 | rs2811464 | rs2811464 | *PLXND1* | 0.0114891 | -0.034 | 1.60E-06 | 2.20E-05 |
| 3 | 170631635 | 170743747 | rs7620985 | rs5402 | *SLC2A2* | 0.0235312 | -0.051 | 2.90E-13 | 1.30E-07 |
| 4* | 68480740 | 68487014 | rs1565070 | rs1565070 | *UBA6* | -0.0105522 | 0.031 | 2.90E-06 | 3.30E-05 |
| 5* | 52100489 | 52122743 | rs59387648 | rs3811978 | *ITGA1* | 0.0150945 | -0.053 | 3.10E-09 | 4.20E-10 |
| 5 | 74352180 | 75042913 | rs2112653 | rs2307111 | *HMGCR-POC5* | -0.0686107 | 0.053 | 1.00E-200 | 3.30E-16 |
| 5 | 78350390 | 78385845 | rs10065798 | rs10944 | *BHMT2* | -0.00840406 | 0.032 | 1.10E-05 | 6.80E-07 |
| 6* | 20396149 | 20396149 | rs79168752 | rs79168752 | *MBOAT1-E2F3* | -0.0158202 | 0.043 | 4.10E-07 | 4.50E-05 |
| 6* | 26200677 | 26338697 | rs6930616 | rs9393698 | *HIST1H4H-BTN3A2* | -0.013272 | -0.036 | 2.10E-11 | 1.30E-06 |
| 6 | 31145920 | 31372569 | rs2523535 | rs887464 | *HLA-C/B* | 0.0130943 | 0.032 | 1.20E-11 | 9.30E-07 |
| 6 | 32127477 | 33514567 | rs9273410 | rs17613292 | *HLA-DQA1/B1* | -0.0199537 | 0.063 | 8.10E-23 | 1.80E-16 |
| 6* | 43804808 | 43821897 | rs68137036 | rs68137036 | *VEGFA-LINC01512* | 0.00943602 | -0.049 | 8.10E-06 | 3.90E-12 |
| 6 | 126629425 | 127167072 | rs9388490 | rs11759026 | *CENPW* | -0.0151964 | -0.067 | 3.20E-15 | 1.30E-18 |
| 7 | 44585052 | 44709372 | rs2008428 | rs79064806 | *NPC1L1-DDX56* | -0.0379635 | -0.049 | 4.50E-63 | 7.10E-06 |
| 7* | 99282772 | 99730628 | rs533394680 | rs535374745 | *over 5 genes* | -0.125826 | -0.37 | 2.70E-07 | 1.50E-06 |
| 7 | 100266081 | 100414386 | rs221793 | rs564449 | *GIGYF1-POP7-EPO* | 0.029001 | -0.048 | 6.70E-20 | 2.20E-06 |
| 8 | 9168146 | 9740317 | rs9987289 | rs17662402 | *PPP1R3B* | -0.0567992 | 0.086 | 1.10E-65 | 9.10E-10 |
| 10 | 64988931 | 65340897 | rs3847326 | rs12245367 | *REEP3* | 0.0124454 | 0.028 | 9.20E-11 | 1.30E-05 |
| 10* | 94809922 | 94809922 | rs117545698 | rs117545698 | *CYP3A4-TRIM4* | 0.0193386 | -0.053 | 6.50E-07 | 4.30E-05 |
| 10 | 101900579 | 102049300 | rs12784396 | rs10883451 | *ERLIN1-CWF19L1* | 0.0124676 | 0.032 | 1.10E-10 | 3.50E-07 |
| 11 | 65422853 | 65604195 | rs4930163 | rs56019505 | *OVOL1-SNX32* | 0.0122885 | 0.041 | 2.50E-06 | 1.20E-05 |
| 12 | 50881148 | 50987378 | rs7316864 | rs34867510 | *DIP2B* | -0.0112604 | -0.029 | 6.70E-08 | 3.20E-05 |
| 12 | 121435778 | 121463140 | rs11065388 | rs12231806 | *HNF1A-OASL* | 0.0178883 | 0.05 | 1.30E-10 | 8.00E-08 |
| 12* | 123762004 | 123891209 | rs34032470 | rs7303754 | *SBNO1* | 0.00955311 | 0.03 | 5.30E-06 | 9.50E-06 |
| 13* | 51067234 | 51099442 | rs9562985 | rs963740 | *DLEU1* | 0.0120389 | 0.039 | 7.40E-08 | 2.60E-08 |
| 15* | 49749735 | 49891094 | rs11639111 | rs11639111 | *FAM227B-FGF7* | -0.0110774 | 0.028 | 1.60E-08 | 2.30E-05 |
| 15 | 62383155 | 62399541 | rs4502156 | rs8037894 | *C2CD4A/B* | -0.00841669 | -0.047 | 1.30E-05 | 3.70E-13 |
| 15* | 63311425 | 63314519 | rs2130165 | rs4774467 | *TPM1* | 0.0111441 | 0.03 | 2.30E-07 | 2.00E-05 |
| 17* | 27557704 | 27637718 | rs56267272 | rs797962 | *NUFIP2* | 0.0137204 | -0.029 | 5.70E-11 | 4.70E-06 |
| 17 | 29628549 | 29749444 | rs7502556 | rs1129506 | *NF1-EVI2A* | -0.0143572 | -0.035 | 2.80E-13 | 8.80E-08 |
| 17 | 46044031 | 46114620 | rs12935890 | rs1076236 | *COPZ2* | -0.0154904 | -0.031 | 6.20E-13 | 2.80E-06 |
| 18* | 55258243 | 55322502 | rs12968116 | rs11660241 | *NARS-ATP8B1* | 0.0200814 | -0.046 | 5.70E-12 | 6.30E-06 |
| 19* | 12176709 | 12193087 | rs73514817 | rs4608438 | *ZNF844* | 0.0115168 | -0.038 | 1.50E-05 | 2.20E-05 |
| 19 | 19320825 | 19793545 | rs2228603 | rs8107974 | *TM6SF2* | -0.107699 | -0.093 | 1.00E-200 | 6.30E-15 |
| 19 | 45324756 | 45428234 | rs147711004 | rs429358 | *APOE* | 0.170228 | 0.08 | 1.00E-200 | 1.80E-18 |
| 19 | 46123929 | 46314504 | rs2238689 | rs9797621 | *GIPR* | 0.016727 | -0.057 | 9.40E-19 | 1.70E-17 |
| 20 | 42958768 | 43042364 | rs1800961 | rs1800961 | *HNF4A* | -0.0755416 | 0.16 | 1.80E-42 | 3.20E-20 |
| 22 | 18415158 | 18486514 | rs11703382 | rs5992928 | *MICAL3* | 0.0111281 | 0.035 | 5.50E-07 | 3.10E-06 |
| 22* | 32367895 | 32410786 | rs74534759 | rs61742196 | *YWHAH-SLC5A1* | -0.0150518 | -0.065 | 2.10E-05 | 2.80E-08 |
| 22 | 44324855 | 44394402 | rs2294915 | rs3747207 | *PNPLA3* | -0.0141373 | 0.049 | 5.70E-10 | 2.10E-10 |

*****indicate loci that were not successfully replicated in GLGC GWAS of LDL-C; bp_min & bp_max: minimum and maximum base pair position for which all SNPs have p<5E-5 for LDL-C and T2D, and with opposite directions of effect. Human build GRCh37/hg19; Beta coefficients correspond to log-odds ratios for T2D, and standard deviations for LDL-C.

**Supplementary Table 4:** Potential functional mechanisms of nearest genes at 31 identified loci.

| **SNP** | **Gene** | **Protein** | **Protein Function** | **References** |
| --- | --- | --- | --- | --- |
| **Fatty Acid Synthesis** | | | | |
| rs5402 | *SLC2A2* | Glucose transporter 2 | Liver specific glucose transporter 2 knockout decreases hepatic glucose uptake, liver triglyceride concentration, and de novo lipogenesis. | {Seyer, 2013 #9} |
| rs649961 | *SLC12A8* | nicotinamide mononucleotide transporter | Transport of nicotinamide mononucleotide, affects hepatic NAD+ concentrations. Given that NAD has a role in glycolysis, TCA cycle, and β-oxidation; NMN transport may affect either lipid synthesis or breakdown. Expression is decreased in humans with NASH. | {Grozio, 2019 #10;Xanthakos, 2015 #11} |
| rs1260326 | *GCKR* | Glucokinase regulatory protein | Normally acts to suppress glucokinase activity by sequestering glucokinase in the nucleus. This sequestration additionally limits glucokinase degradation and allows for rapid mobilizaton of glucokinase activity from the nucleus when glucose is high. In turn, GCKR knockout decreases hepatic glucokinase protein and activity, resulting in impaired glucose clearance when blood glucose is high. This depressed hepatic glucose clearance would be expected to limit carbon flux toward fatty acid synthesis. In fact, this exact variant in GCRP is associated with fatty liver. | {Farrelly, 1999 #7;Grimsby, 2000 #6;Santoro, 2012 #8} |
| rs12231806 | *HNF1A-OASL* | Hepatocyte nuclear factor 1 alpha | [Chronic high fat diet exposure decreases hepatic HNF1A mRNA expression, while deletion of hepatic HNF1A increases hepatic lipid concentration. This is due to increased hepatic *de novo* lipogenesis.](https://www.jci.org/articles/view/121994%20%20beta%20cell%20dysfunction) | {Ni, 2017 #50} |
| **Hepatic Lipid Uptake** | | | | |
| rs79064806 | *NPC1L1-DDX56* | NPC1 Like Intracellular Cholesterol Transporter 1 | Found on the brush border of enterocytes and the canalicular membrane in hepatocytes where it is involved in dietary cholesteral absorption and re-uptake of cholesterol from bile. NPC1L1 knockout reduces cholesterol absorption in mice, while expression of NPC1L1 in mouse livers increases cholesterol reabsorption from bile. Ezetimibe, a cholesterol lowering drug blocks NPC1L1 activity. | {Davis, 2004 #13;Tang, 2011 #15;Wang, 2018 #14} |
| rs41279732 | *SORT1-PSMA5* | sortilin 1 | KO decreases NPC1 Like Intracellular Cholesterol Transporter 1 expression, decreases adipose tissue mass. SORT1 is highly expressed in adipocytes, facilitates formation and export of VLDL from liver. | {Hagita, 2018 #1; Chen, 2019 #56} |
| rs429358 | *APOE* | Apolipoprotein E | Apoliprotein E knockout mouse is hypercholesterolemic due to poor lipoprotein clearance, yet resistant to high fat diet induced hepatic triglyceride accumulation. In insulin resistant MKR mice APOE knockout improves glucose clearance and insulin sensitivity. | {Karavia, 2011 #16;Kawashima, 2009 #17} |
| rs2307111 | *HMGCR-POC5* | HMGCoAReductase | HMGCR is a key regulatory enzyme in cholesterol synthesis, directly affecting circulating LDL concentrations. Inhibition of HMGCR in the liver stimulates the LDL-receptors, which results in an increased clearance of LDL from the bloodstream and a decrease in blood cholesterol levels | {Jiang, 2018 #19;Pocathikorn, 2010 #20} |
| **Lipid Export** | | | | |
| rs8107974 | *TM6SF2* | transmembrane 6 superfamily member 2 | rs58542926 has previously been associated with increased hepatic triglyceride concentrations and altered incidence of hepatic fibrosis. Inactivation of TM6SF2 prevents lipid loading of VLDL, but does not prevent apolipoprotein B100 secretion. | {Liu, 2014 #21;Smagris, 2016 #22} |
| rs9797621 | *GIPR* | glucose-dependent insulinotropic polypeptide receptor | GIPR knockout in adipose tissue decreases high fat diet induced weight gain, hepatic steatosis, insulin reistance, and liver mass, without altering adipose tissue mass. Visceral adipose tissue mass is affected by variants in GIPR and an antibody against GIPR has been shown to decrease body mass, serum insulin, and serum triglycerides. | {Joo, 2017 #23;Wang, 2017 #24;Killion, 2018 #25} |
| rs1800961 | *HNF4A* | Hepatocyte Nuclear Factor 4 Alpha | Knockdown of HNF4A results in development of fatty liver, a plasma triglycerides, total cholesterol, and high-density lipoprotein cholesterol. The decreae in lipids may be due to decreased lipogenesis and VLDL production, as HNFA knockdown decreased fatty acid synthase, hydroxy methyl glutaryl CoA reductase, low density lipoprotein receptor, and Apolipoprotein B mRNA expression. | {Yin, 2011 #27} |
| rs10883451 | *ERLIN1-CHUK-CWF19L1* | Endoplasmic Reticulum Lipid Raft-Associated Protein 1 - Component Of Inhibitor Of Nuclear Factor Kappa B Kinase Complex - CWF19 Like Cell Cycle Control Factor 1 | This gene cluster has previosly been shown to induce fatty liver. In fact, this exact SNP was previously identified. ERLIN1 is associated with degradation of IP3 receptors, CHUK inhibits nuclear factor kappa B kinase complex, and CWF19 Like Cell Cycle Control Factor 1 is associated with lean body mass. | {Feitosa, 2013 #28} |
| rs2741042 | *UGT1A8* | UDP Glucuronosyltransferase Family 1 | UGT1A8 is Involved in adding glucoronic acid (a sugar) to lipophillic molecules so that they can be transported in water based solvents. UGT1A8 is expressed predominantly in intestinal cells, but is upregulated by poly-unsatturated fatty acids in hepatocytes. UGT1A8 is likely to be involved in cholesterol absorption at the small intestine and excretion in bile. | {Ziegler, 2015 #12} |
| **Peripheral Tissue Lipid Balance** | | | | |
| rs1129506 | *NF1-EVI2A* | neurofibromin | Skeletal muscle NF1 deletion increases skeletal muscle lipid concentration in mice. Patients with neurofibromatosis type 1, a result of NF1 mutations, have lower fasting blood glucose and appear to have improved insulin sensitivity. | {Sullivan, 2014 #31;Martins, 2016 #32;Martins, 2018 #33} |
| rs34867510 | *DIP2B* | DIP2 disco-interacting protein 2 homolog B | decreased serum FFAs |  |
| rs17036170 | *PPARG* | PPARy | PPARγ activity in adipose tissue encourages free fatty acid uptake and differentiation preventing the lipotoxicity that can accompany storage of fatty acids in tissues other than adipose tissue. The thiazolidinediones, commonly used to treat type II diabetes, are PPARγ agonists. PPARγ activity in the liver encourages hepatic lipid storage and an adipocyte like phenotype. | {Hammarstedt, 2005 #34;Medina-Gomez, 2007 #35;Wolf Greenstein, 2017 #36} |
| rs17613292 | *HLA-DQA1/B1* | major histocompatibility complex, class II, DQ alpha 1 | Inflammation is a key feature of type 2 diabetes mellitus and MHC Class II proteins play an essential role in adipose tissue inflammation, the associated increase in adipose tissue lipolytic flux and the metabolic dysfunction in obesity. | {Deng, 2013 #45} |
| **Fatty Liver of Unknown Origin** | | | | |
| rs12245367 | *REEP3* | RECEPTOR EXPRESSION-ENHANCING PROTEIN 3 | total Cholesterol | {Chalasani, 2010 #3} |
| rs17662402 | *PPP1R3B* | Protein Phosphatase 1 Regulatory Subunit 3B | PPAR1R3B regulates hepatic glycogen synthesis. Knockout of PPAR1R3B decreases hepatic glycogen storage and overexpression of PPAR1R3B increased hepatic glycogen storage. Decreased glycogen storage promotes hepatic gluconeogenesis, ketogenesis, and lipid β-oxidation to meet the glucose and energy needs of non-hepatic tissues. | {Mehta, 2017 #37} |
| rs3747207 | *PNPLA3* | Patatin-like phospholipase domain-containing protein 3 (PNPLA3), adiponutrin (ADPN), acylglycerol O-acyltransferase or calcium-independent phospholipase A2-epsilon (iPLA2-epsilon) | This enzyme catalyzes the synthesis of phosphatidic acid a key step in triglyceride synthesis. In addition, PNPLA3, expressed in both cytosol and mitochondria plays a key role in triglyceride hydrolysis. In humans, sequence variants have been established to affect fatty liver. | {Jenkins, 2004 #38;He, 2010 #39;Kumari, 2012 #40} |
| rs56019505 | *OVOL1-SNX32* | Ovo like transcriptional repressor 1-Sortin Nexin 32 | Sortin nexin 32 is a HNF4A target gene that docks on the membrane by binding phosphoinositol and regulates membrane trafficking through the endocytic pathway. Although no specific findings link SNX32 to specific receptors, this key protein in the cargo selection complex could alter trafficking of a number of cell surface receptors that alter lipid metabolism (LDL-R, Insulin receptor, glucagon receptor). | {Mathelier, 2014 #41;Sandelin, 2004 #42;Seet, 2006 #43;Naslavsky, 2018 #44} |
| rs887464 | *HLA-C/B* | major histocompatibility complex, class I, C and B | HLA-C and HLA-B alleles affect the incidence of obesity and both HLA-C and HLA-B affect the fibrotic response to NAFLD and progression to NASH | {Shen, 2018 #46;Karrar, 2019 #47} |
| **Insulin Secretion** | | | | |
| rs8037894 | *C2CD4A/B* | C2 Calcium Dependent Domain Containing 4A | C2CD4A/B variants affect GSIS in humans. In fact, the rs7172432 variant impairs glucose-stimulated insulin response. Knockout of CDC4A in the mouse results in in β-cell dysfunction and a lack of glucose stimulated serum insulin release. A common variant increases the risk of type 2 diabetes. | {Grarup, 2011 #4;Kuo, 2019 #5;Kycia, 2018 #60} |
| rs5402 | *SLC2A2* | Glucose transporter 2 (GLUT2) | Beta-cells express GLUT2 and the influx of glucose into the β-cell is essential for the insulin secretory response to glucose. | {Guillam, 2000 #2} |
| rs5992928 | *MICAL3* | Molecule Interacting with CasL | MICAL3 is involved in exocytosis of secretory vesicles. In turn, changes in MICAL3 expression or function may alter insulin secretion. Experimental mutation of MICAL3, inactivating the monoonxygenase domain prevented vesicular fusion and led to a build up of secretory vesicles within the cell. | {Grigoriev, 2011 #48} |
| rs12231806 | *HNF1A-OASL* | Hepatocyte nuclear factor 1 alpha | [HNF1A mutation can prevent glucose stimulated insulin release from β-cells by altering gene expression of enzymes in glucose metabolism and vesicular exocytosis](https://www.jci.org/articles/view/121994%20%20beta%20cell%20dysfunction) | {Haliyur, 2019 #49} |
| rs9797621 | *GIPR* | Glucose-dependent insulinotropic polypeptide receptor | GIP signaling through the GIPR stimulates β-cell insulin release. | {Fujita, 2010 #51;Harada, 2017 #52} |
| **Insulin Action/Peripheral Glucose Uptake** | | | | |
| rs12144658 | *FNDC7-STXBP3* | Syntaxin Binding Protein 3 (also known as Munc 18-3 or Munc 18c) | Together with STX4 and VAMP2, may play a role in insulin-dependent movement of GLUT4 and in docking/fusion of intracellular GLUT4-containing vesicles with the cell surface in adipocytes.  In fact, insulin triggers the formation of a PKCζ–80K-H–STXBP3 complex that increases GLUT4 translocation to the plasma membrane. | {Hodgkinson, 2005 #54;Khan, 2001 #55;Thurmond, 1998 #56} |
| rs564449 | *GIGYF1-POP7-EPO* | GRB10 Interacting GYF Protein 1 | GIGYFI interacts with GRB10 to enhance insulin like factor-1 (IGF-1) receptor signalling. Since GRB10 acts to inhibit both insulin receptor and IGF-1 receptor signalling, GIGYF1 may also enhance insulin receptor signaling. At the β-cell this may increase insulin release, while at other tissues it may increase glucose disposal. | {Giovannone, 2003 #57;Li, 2013 #58;Wang, 2007 #59} |
| **Unknown Action - Unclassified** | | | | |
| rs10944 | *BHMT2* | Betaine--Homocysteine S-Methyltransferase 2 |  |  |
| rs10460461 | *LINC01101-GLI2* | Long intergenic non-protein coding RNA 1101. GLI2 |  |  |
| rs1076236 | *COPZ2* | Coatomer Protein Complex Subunit Zeta 2 |  |  |
| rs116097055 | *THADA* | Thyroid Adenoma-Associated Gene |  |  |
| rs11759026 | *CENPW* | Centromere Protein W |  |  |

**Supplementary Figure 1:** Sex-stratified T2D prevalence by LDL-C decile. T2D prevalence is shown as a percentage with error bars corresponding to the Clopper-Pearson confidence interval.

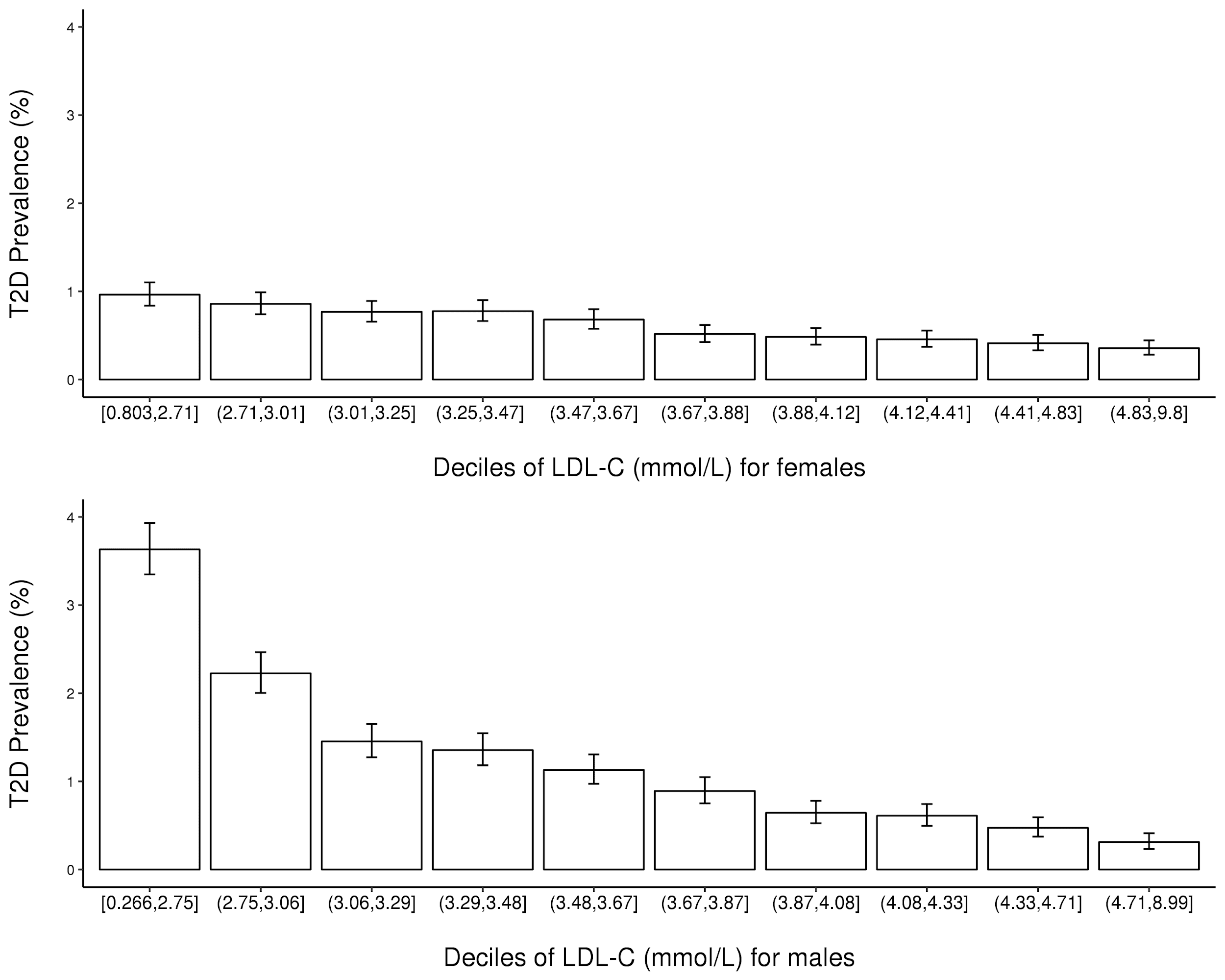

**Supplementary Figure 2:** Age-stratified T2D prevalence by LDL-C decile. T2D prevalence is shown as a percentage with error bars corresponding to the Clopper-Pearson confidence interval.

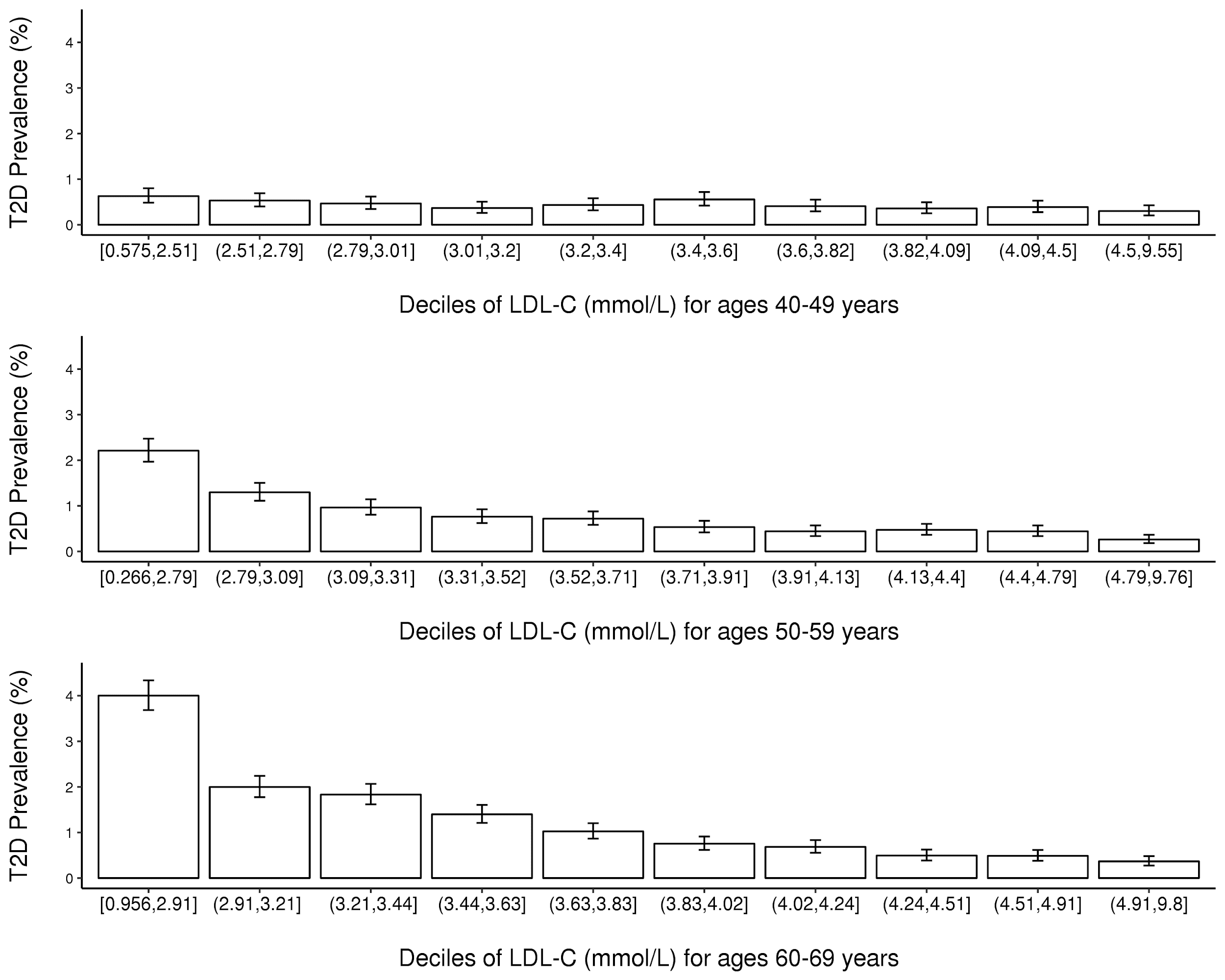

**Supplementary Figure 3:** T2D prevalence by TG decile. T2D prevalence is shown as a percentage with error bars corresponding to the Clopper-Pearson confidence interval.

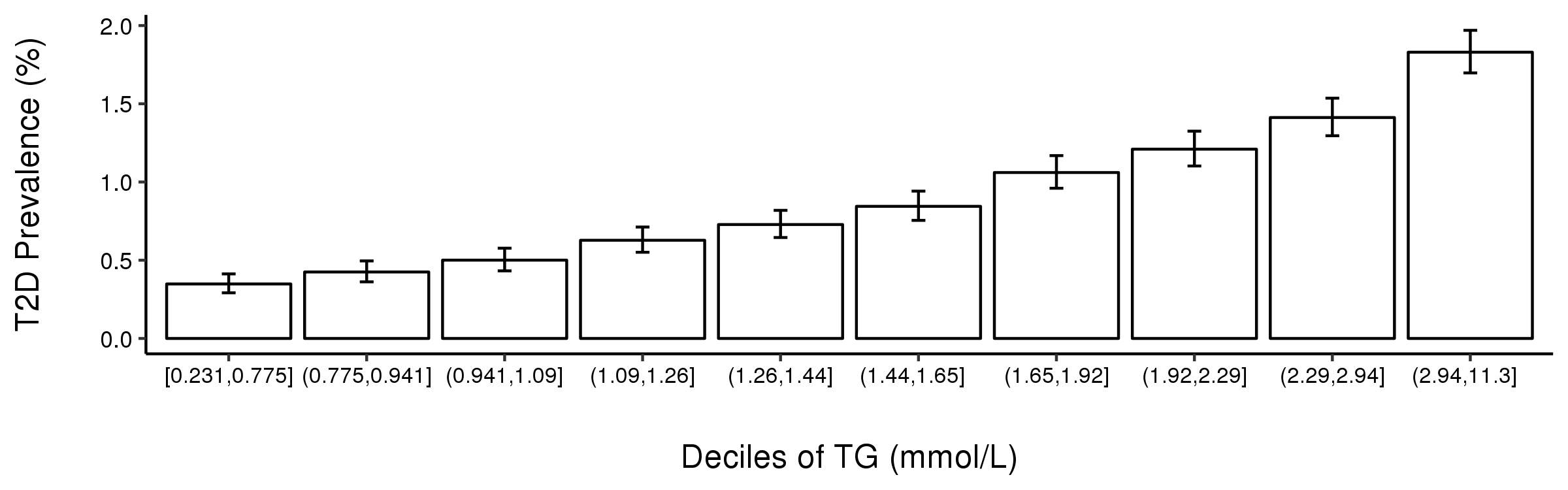

**Supplementary Figure 4:** Mendelian randomization scatter plot.

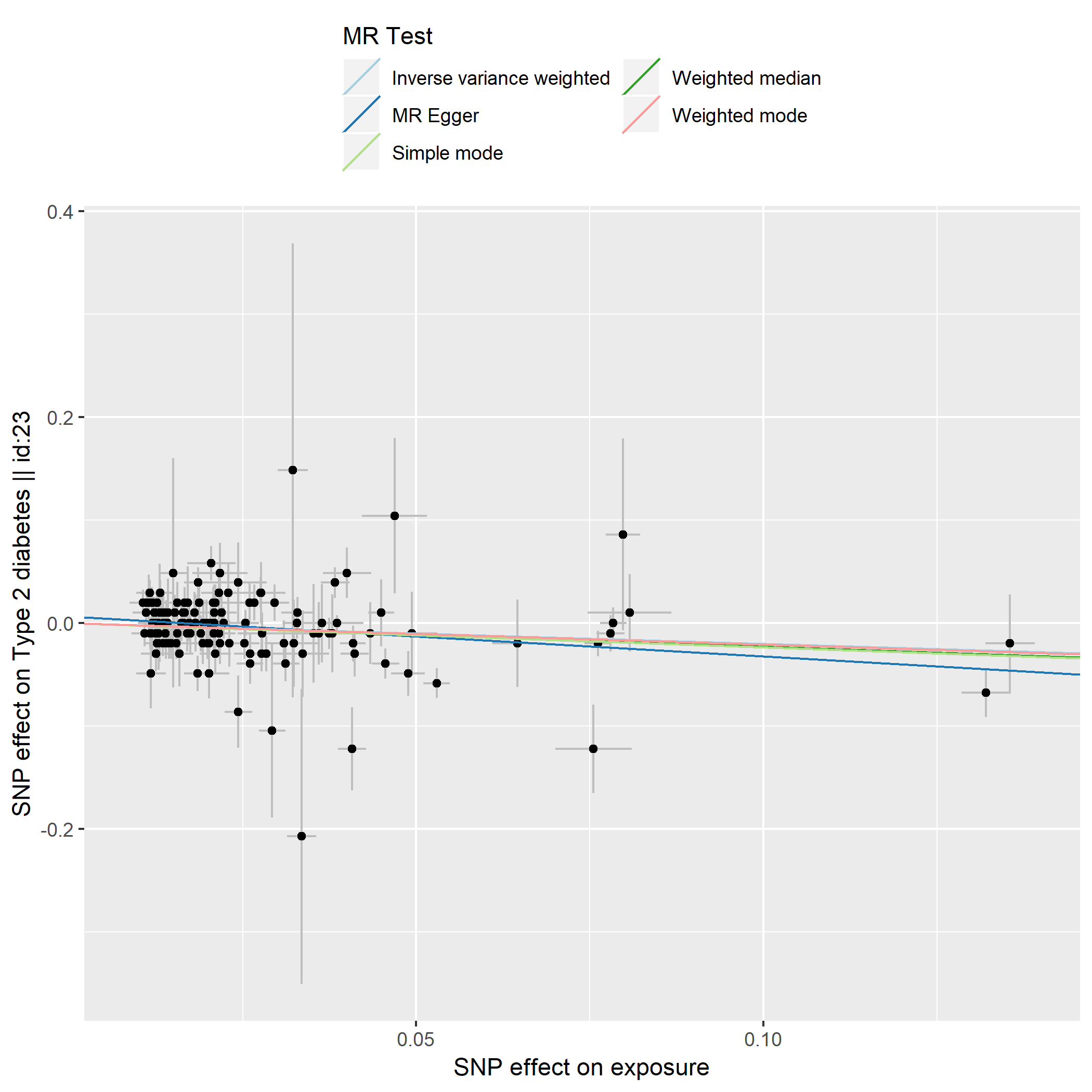

**Supplementary Figure 5:** Forest plot from Mendelian randomization of LDL-associated SNPs on T2D outcome.

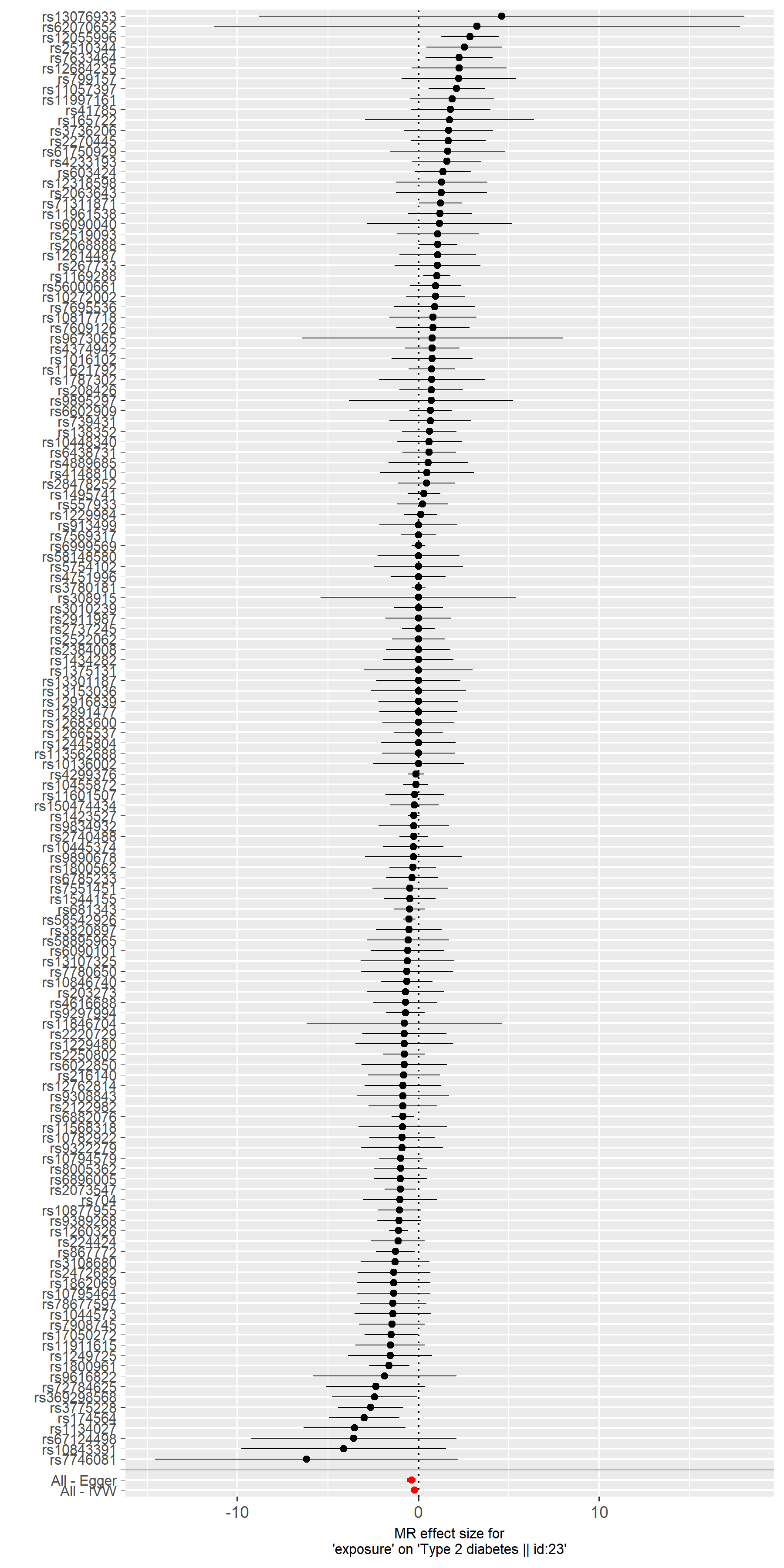

**Supplementary Figure 6:** Leave-one-out plot from Mendelian randomization of LDL-associated SNPs on T2D outcome.

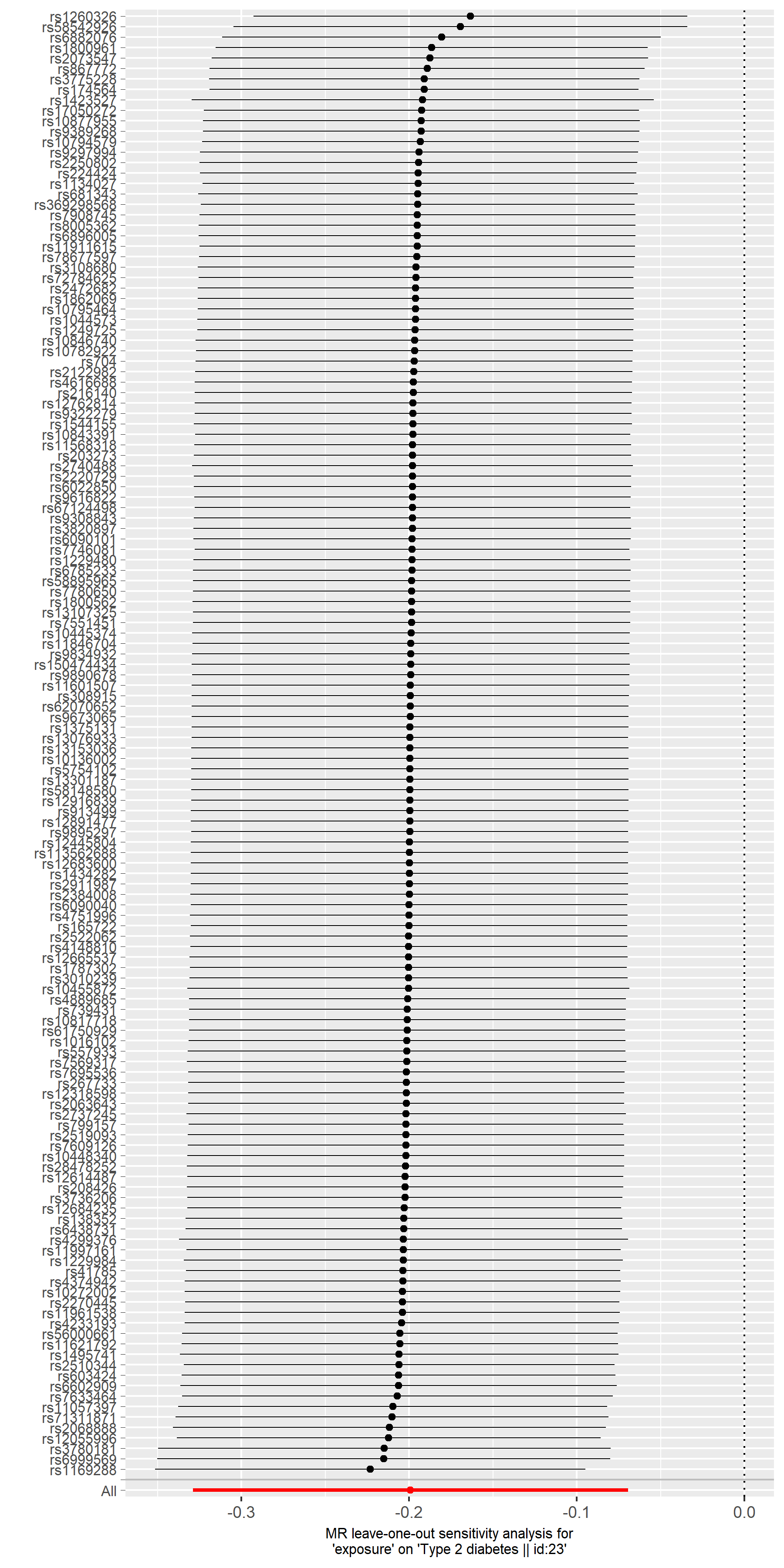
